# Exploring the Sequence Fitness Landscape of a Bridge Between Protein Folds

**DOI:** 10.1101/2020.05.20.106278

**Authors:** Pengfei Tian, Robert B. Best

## Abstract

Most foldable protein sequences adopt only a single native fold. Recent protein design studies have, however, created protein sequences which fold into different structures apon changes of environment, or single point mutation, the best characterized example being the switch between the folds of the GA and GB binding domains of streptococcal protein G. To obtain further insight into the design of sequences which can switch folds, we have used a computational model for the fitness landscape of a single fold, built from the observed sequence variation of protein homologues. We have recently shown that such coevolutionary models can be used to design novel foldable sequences. By appropriately combining two of these models to describe the joint fitness landscape of GA and GB, we are able to describe the propensity of a given sequence for each of the two folds. We have successfully tested the combined model against the known series of designed GA/GB hybrids. Using Monte Carlo simulations on this landscape, we are able to identify pathways of mutations connecting the two folds. In the absence of a requirement for domain stability, the most frequent paths go via sequences in which neither domain is stably folded, reminiscent of the propensity for certain intrinsically disordered proteins to fold into different structures according to context. Even if the folded state is required to be stable, we find that there is nonetheless still a wide range of sequences which are close to the transition region and therefore likely fold switches, consistent with recent estimates that fold switching may be more widespread than had been thought.

**Author Summary:** While most proteins self-assemble (or “fold”) to a unique three-dimensional structure, a few have been identified that can fold into two distinct structures. These so-called “metamorphic” proteins that can switch folds have attracted a lot of recent interest, and it has been suggested that they may be much more widespread than currently appreciated. We have developed a computational model that captures the propensity of a given protein sequence to fold into either one of two specific structures (GA and GB), in order to investigate which sequences are able to fold to both GA and GB (“switch sequences”), versus just one of them. Our model predicts that there is a large number of switch sequences that could fold into both structures, but also that the most likely such sequences are those for which the folded structures have low stability, in agreement with available experimental data. This also suggests that intrinsically disordered proteins which can fold into different structures on binding may provide an evolutionary path in sequence space between protein folds.

## Introduction

There is an enormous variety of protein sequences found in nature, with around 170 million non-redundant sequences registered in the Refseq database [1] at the time of writing. A significant fraction of these, approximately 1/3 in eukaryotes [2, 3], are intrinsically disordered. The sequence diversity of the the remainder, which fold to a specific structure, belies a simplicity in the structures to which they fold: most folded proteins can be classified into one or more independently folding units, or domains [4], and the number of domains which have a distinct structure, numbering in the thousands, is much more limited than the number of sequences that fold to these structures [5, 6]. Here, by distinct structure, we mean proteins which have the same overall fold, i.e. that the three dimensional arrangement of the backbone and secondary structure elements is similar. While the number of experimentally determined structures in the protein data bank continues to grow rapidly, the number of known folds is increasing only very slowly, suggesting that most existing naturally occurring folds are already known [6].

Recent advances in protein design have also shown it is possible to design completely novel folds, not observed in nature [7]. Therefore the number of folds sampled by evolution is smaller than the number possible. Indeed a molecular simulation study exploring possible protein architectures hinted that the number of possible folds may even be considerably larger than those currently known [8]. These results, as well as bioinformatics analysis [9], suggest that the emergence of new folds is a very rare event in protein evolution. How, then, do new folds arise? One possible route is via evolution of existing ones [10, 11]. In this scenario, there would be pathways in sequence space between the two folds, in which the intermediate sequences would have some propensity to fold into both structures. Such sequences are expected to be very rare, given that the fraction of possible random sequences which actually fold to a specific, stable backbone structure is already extremely tiny [12–16].

Remarkably, however, there are several naturally occurring examples in which the same protein sequence can adopt two completely different stable folds apon changes in conditions [17], for example changes in pH (lymphotactin [18]), or binding to another molecule (KaiB [19]). It has also been possible to design proteins which can switch folds: a temperature-sensitive local switch of structure between helix and sheet was obtained in a designed version of arc-repressor [20, 21], and more recently sequences have been designed which make the dramatic switch between the all-*α* GA and *α*/*β* GB folds of streptococcal protein G apon single-point mutation, or addition of a binding partner [17, 22]. These so-called “metamorphic” proteins [23] have sparked interest for their biophysical properties, their potential roles as molecular switches, as well as their possible link to protein evolution. Bioinformatics analysis has suggested that such fold switches may be even more widespread than currently thought [24, 25].

The designed fold switch between the all-*α* GA and the *α/β* GB folds is the best experimentally characterized metamorphic protein pair (Fig. 1). Via a systematic, and conservative, alteration of the sequence, Bryan, Orban and co-workers have demonstrated that it is possible to switch the structure of the GA domain (Fig. 1 green) to the GB domain (Fig. 1 purple) [26]. In some cases, a single point mutation is enough to switch from one structure to another, and some variants appear to be able to populate both structures, under different conditions [22]. The rich structure and stability data describing a mutational pathway between the GA and GB folds has inspired a number of theoretical studies of the fold switching phenomenon [27]. The models used in such studies are, by necessity, usually highly simplified: for example, a reduced three-letter protein model was used to study the sharp fold switch caused by a short mutational path [28]. 2-D lattice models can also be used as generic models to explore the general features of sequences that act like evolutionary bridges [29, 30]. The above models attempt to model both the changes in sequence space, as well as the actual folding of the chain in three dimensions. This requirement necessarily limits them to model systems (reduced alphabets, lattice models). In order to describe and predict protein sequences which act as a bridge between the specific GA and GB folds, a more detailed model is needed. One approach is to use all-atom physical force fields [31, 32], but these are very computationally expensive and still not fully predictive. By combining an all-atom physical force field with an additional energy term for native contacts it was possible to determine the free energy differences between fold switch mutants [33]. However, adjusting the relative weight between the native contacts energy and physical energy is not trivial, and the application is limited to a few mutants due to the computational cost involved. Some sequence-dependent models have been parametrized to fit the fold propensity of the mutations at the interface of the GA/GB fold, but the overall landscape of the bridge between two folds was not characterized [32, 34].

**Figure 1.**
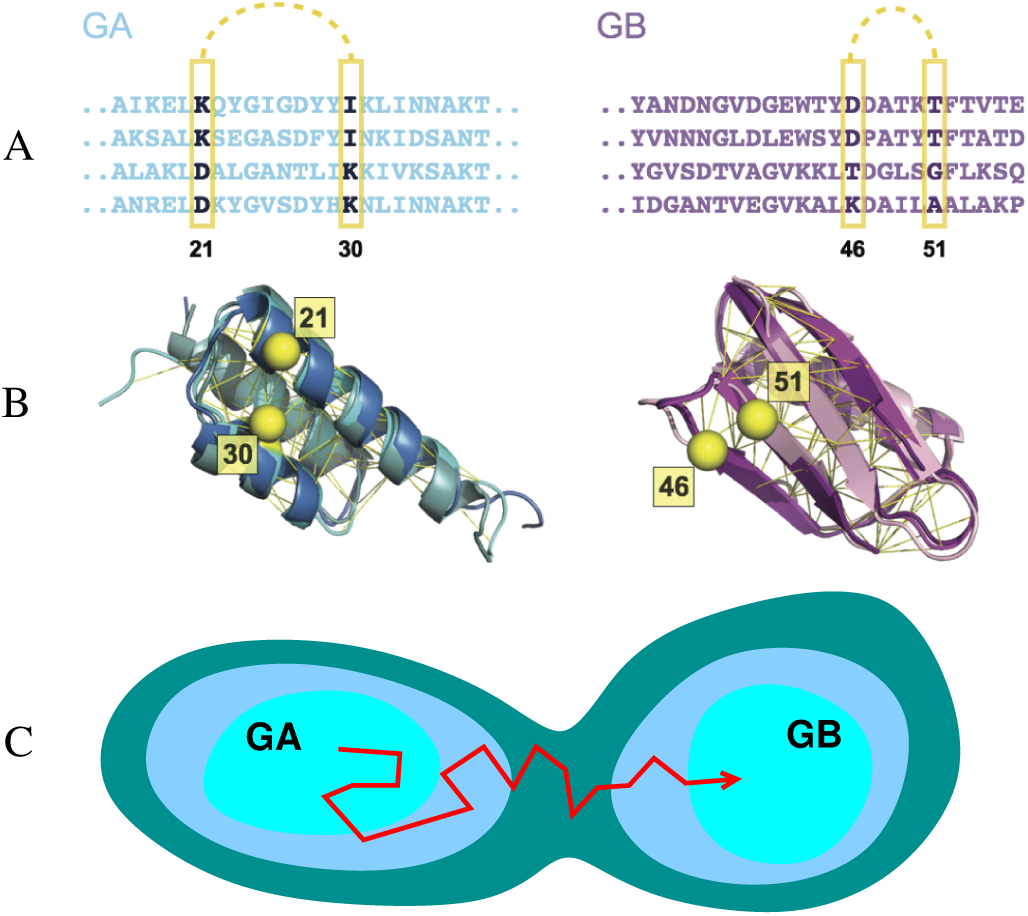
Sequence-based models for the GA and GB domains of streptococcal protein G. Many sequences (A) fold to each structure (B): e.g. structures of three naturally occurring sequences with the GA fold (pdb ID 2fs1, 1gjs and 2j5y) and three with the GB fold (pdb ID 1pga, 2lum and 1igd) are shown on the left and right respectively. Contacts between pairs of residues in the native structure (C*β* atoms of example pairs in yellow) impose mutual constraints on the types of residues which can occupy these positions in the sequence alignment. For instance, strong covariance are detected between the amino acid pair at residue 21 and 30 for GA sequences and between residue 46 and 51 for GB sequences. The C*β* atom of these residues are illustrated in yellow sphere. The UniProtKB ID of these example sequences for GA are Q51918 FINMA, G5KGV3 9STRE, G5K7M6 9STRE and Q56192 STAXY. And the ones for GB are SPG1 STRSG, E4KPW8 9LACT, F9P4J6 STRCV and G5JZF8 9STRE. (C) Simple model for the emergence of new folds via evolutionary drift in sequence space between basins of attraction corresponding to the GA and GB domains.

Our goal in this work was to develop a model for a sequence-space fitness landscape representing the joint fitness for the GA and GB folds, and to characterize pathways in sequence space between folds (Fig. 1C). We use as input the observed sequence variation of protein homologues, which captures the covariation of amino acids at different sites; previously, we have demonstrated that it is possible to use such models to predict the effects of mutations for the proteins we have considered [13], as well as other those from previous studies [35–38]. We have even shown that it is possible to use such models to design novel sequences that fold stably into either a GA, GB or SH3 fold, representing the three basic classes of protein structure (all-*α, α/β*, all-*β* respectively) [39]. Here, we generalize such coevolutionary models to allow for transitions between the basins of attraction in sequence space corresponding to each fold. By using Monte Carlo simulations to sample transitions between these basins, we have described the characteristics of the mutational bridge between the GA and GB folds in sequence space. The rapid exploration of sequence space made possible with such a model allows us to investigate the effect that different requirements on the protein stabilities have on evolutionary dynamics [40, 41].

## Results

### Statistical Model of GA and GB Sequences

Maintaining the structure of the folded state is an important constraint on natural selection in protein evolution [35, 42, 43]. Therefore, proteins from the same family, which share the same fold, should contain common features in their sequences, both in the propensities of residues to be at certain positions, as well as the covariation between different sites which are in contact in the native state. The variation of the related sequences contained in a multiple sequence alignment (MSA) contains rich evolutionary information about structural and functional constraints (Fig. 1).

In our work, we have built a model for the fitness of a given sequence to fold into a given structure, based on the covariation of sequences sampled in nature. The model for each protein family is parameterized using residue-residue coevolutionary information, which has previously been used to predict native contacts of protein structures [44–48], protein-protein interations [49–51] and RNA structures [52, 53]. Firstly, as shown in the MSA fragment in Fig. 1, there is a propensity for certain residues to be found at a given position of the sequence. Secondly, there are correlations between the propensity at different sites, i.e. if one residue mutates, the proximal residues in the three dimensional structure will also likely mutate to maintain compatible physical and chemical interactions [54] (e.g. having Asp at position 21 and Lys at position 30 is favourable, but if position 21 is changed to Lys, it is unfavourable to have Lys at position 30). These propensities are approximated by the following Potts-like likelihood function *P* (*A*_1_, *A*_2_, .., *A*_*L*_), representing the likelihood of a given amino acid sequence *A*_1_, *A*_2_, .., *A*_*L*_, of length *L* for a particular protein fold,

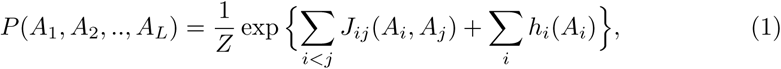

In this function, the parameters *h*_*i*_ represent the single-site propensities for a given amino acid *A*_*i*_ at position *i*, while *J*_*ij*_ represents the propensity for amino acids *A*_*i*_ and *A*_*j*_ to be at positions *i* and *j*. These parameters are optimized to be consistent with the sequences observed in the MSA, using a pseudolikelihood optimization scheme [55]. From this probability, we associate an energy (“evolutionary Hamiltonian”) with a given sequence *x*, via *E*_EH_(*x*) = − ln *P* (*x*) (in units of *k*_B_*T*). We have built such a model for both protein families GA and GB. *E*_EH,GA_ and *E*_EH,GB_ are the two Hamiltonians inferred from the homologous sequences of GA and GB respectively using Equation 1; in earlier work, we showed that it was possible to design stably folded proteins using such evolutionary energy functions, for each of GA, GB and SH3 domains [39]. We first verified that Metropolis Monte Carlo simulations using the evolutionary energies *E*_GA_ or *E*_GB_ can recapitulate both the energy distribution of the sequences from the MSA of GA or GB (S1 Text Fig. A) as well as the amino acid composition frequencies (S1 Text Fig. B).

Some properties of the potentials are illustrated in Fig. 2. As expected, the sequences used to build the model occupy the lowest energy region in each case (Fig. 2A,B). The synthetic sequences designed by Bryan and co-workers (S1 Table A) [22, 56–58] can be divided into those which are unstable, which have the highest energy with either *E*_GA_ or *E*_GB_, and those which fold to either GA or GB, which have energies intermediate between the respective training set and those that do not fold. We have also calculated the energies of sequences which we have generated by selecting at random from the residues which occur at each position in the sequence alignment, i.e. with no energy bias (grey histogram in Fig. 2A,B). It is clear that the unstable designed sequences still have a significant propensity for the target fold, since their energies are much closer to the stable designed sequences than to random sequences. In Fig. 2C,D, we compare the folding midpoint temperature *T*_*m*_ (data in S1 Table B), a measure of folded state stability, and the statistical energy for each sequence. We observe a good correlation in each case (rank correlation coefficients of 0.86 and 0.92 for GA and GB respectively), with the unstable sequences also having the highest statistical energy. Such a correlation is expected if protein stability is an important consideration for natural selection, and has been observed also for other proteins [35–37]. In S1 Text Fig. C we show that a similar correlation exists with folding free energies, where those are available.

**Figure 2.**
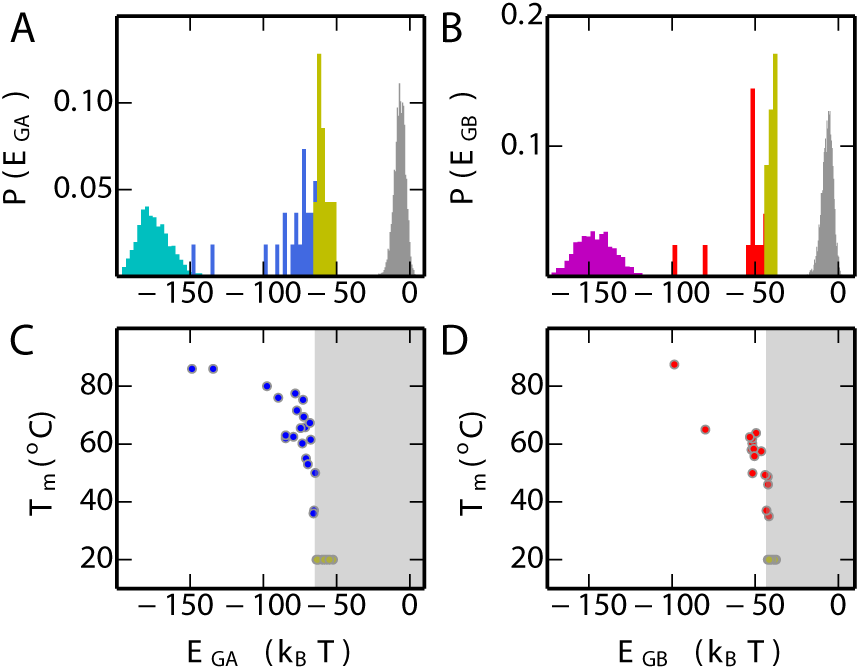
Properties of the single-fold models. (A) Distribution of *E*_GA_ for the GA homologs used to parameterize *E*_GA_ (cyan), synthetic sequences which are dominated by GA fold (blue) state in equilibrium, unstable synthetic sequences (yellow) and randomly generated sequences (grey). (B) Distribution of *E*_GB_ for the GB homologs used to parameterize *E*_GB_ (purple), synthetic sequences which are dominated by GB fold (red) state in equilibrium, unstable synthetic sequences (yellow) and random sequences (grey). (C) The correlation between the folding temperature (*T*_*m*_) and *E*_GA_ for synthetic sequences of GA. Stable mutants are blue symbols, unstable are yellow symbols with *T*_*m*_ set to 20°C for plotting purposes. (D) The correlation between *T*_*m*_ and *E*_GB_ for experimental mutants of GB (stable: red, unstable: yellow, *T*_*m*_ set to 20°C).

### A Combined Fitness Landscape for Two Protein Folds

The models for GA and GB separately describe the fitness of sequences for each fold. In order to realize our goal of studying transitions between sequences which fold into GA and those which fold into GB, we require a single energy surface. A natural way to achieve this is to add the individual likelihood functions exp[−*E*_GA_] and exp[−*E*_GB_] or to use the more general combined energy function *E*_comb_ defined for sequence *x* as [59],

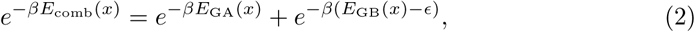

where *β* is the inverse of a “mixing temperature” *T*_mix_ that determines the extent of mixing between the two potentials and is fixed here to 1.0. *ϵ* is an energy offset which sets the relative free energy of the two basins. Sequences from the GA MSA and GB MSA occupy the two minima of *E*_comb_, with the sequences near the barrier of the combined potential *E*_comb_ being putative “bridges” between the two folds.

There is only one undetermined parameter in the combined energy function *E*_comb_, i.e. the offset energy *ϵ*. We find an appropriate value for *ϵ* using the committor function *ϕ*_A_(*x*) [60–62], defined as the probability that trial Monte Carlo simulations in sequence space (described in more detail below), initiated from sequence *x*, first reach the free energy minimum corresponding to the GA fold rather than GB: ideally sequences which are known to fold to GA should lie within the basin of attraction of GA in sequence space and have *ϕ*_A_ > 0.5, and those folding to GB would have *ϕ*_A_ < 0.5. An optimal *ϵ* = 23.0 is chosen for which the known propensity of a given sequence for the GA (versus GB) fold is correlated with the splitting probability *ϕ*_*A*_. With this choice, we find that *ϕ*_*A*_ is a good predictor of the favoured fold. Most of the designed sequences, such as GA30, GB30, GA77, GB77, GA88 and GB88, only ever populate one fold in experiment: Consistent with that, the *ϕ*_*A*_ estimated for these sequences is very close to 1.0 or 0.0. On the other hand, the mutants GA98, GB98, GB98-T25I and GB98-T25I/L20A all can adopt both GA and GB folds, either at equilibrium, or in the presence of binding partners. The GA fold is the most populated in the GA98 and GB98-T25I mutants, with a small population of the GB fold, ∼5% for GB98-T25I and ∼1% in GA98 [22]. For the GB98 and GB98-T25I/L20A mutants, the major population is the GB fold. The minor GA population in GB98 is larger than in GB98-T25I/L20A, although the exact populations have not been determined [22, 26]. The *ϕ*_*A*_ values of these four mutations in the Fig. 3A, reproduce these observations, with *ϕ*_*A*_(GA98) > *ϕ*_*A*_(GB98-T25I) > 0.5 > *ϕ*_*A*_(GB98) > *ϕ*_*A*_(GB98-T25I/L20A). Note that alternative choices of *ϵ* will shift the position of the fold interface (i.e. *ϕ* = 0.5) while the relative ranking of *ϕ* over the different mutants is not changed (S1 Text Fig. D).

**Figure 3.**
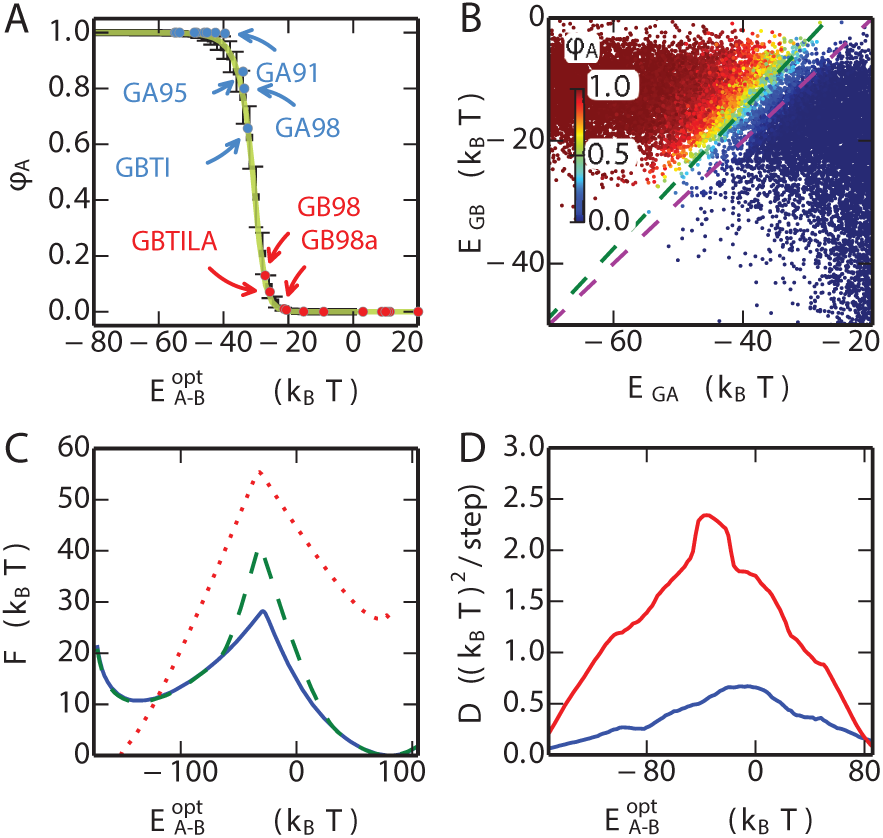
One-dimensional energy landscape capturing fold switch. (A) The committor for reaching the GA fold, *ϕ*_*A*_ is plotted for the experimentally characterized mutant sequences with blue (GA fold) and red (GB fold) symbols. The mean and standard deviation of *ϕ*_*A*_ for an equilibrium sample of sequences at given values of the optimized coordinate 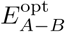 are shown by black symbols and errorbars. The theoretical committor from a 1D diffusion model is shown in yellow. (B) The *ϕ*_*A*_ values (colours) are projected onto *E*_GA_ and *E*_GB_ for each sequence. Purple and blue broken lines are perpendicular to the original coordinate *E*_*A*−*B*_ = *E*_GA_ − *E*_GB_ and the optimized coordinate 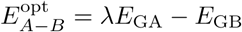 respectively (*λ* = 1.13). (C) Free energy profile of the combined model for the natural mutations (blue), natural mutations with stability constraints (green) and the binary mutations (red). (D) The profile of position-dependent diffusion coefficients for the natural mutations (blue) and the binary mutations (red).

### Exploring Fold Switching in Sequence Space

Guided by the combined model *E*_comb_, we have explored the joint fitness landscape of the the two folds by the Monte Carlo simulation, in which a Metropolis criterion is used to accept or reject trial moves in sequence space. Such simulations correspond to a highly simplified model of protein evolution. We consider two different move sets in our simulations: “natural” and “binary” mutations. For natural mutations a new residue type is chosen with equal probability from those amino acids which are found at that position in the MSA of GA and GB. This restriction is made to avoid exploring regions of sequence space about which our statistical potential has no information and would therefore not be reliable. In the more conservative binary mutation scheme, the only allowed residues are those found in the reference GA and GB sequences (all of the sequences designed by Bryan et al. fall within this scheme [57]).

In order to characterize the fitness landscape, including regions with low population, we initially performed umbrella sampling using as reaction coordinate the energy gap *E*_A-B_(*x*) = *E*_GA_(*x*) − *E*_GB_(*x*), which has proved a useful coordinate in the context of previous problems involving mixed energy functions [63, 64]. This coordinate also separates quite well the sequences folding into GA vs GB (S1 Text Fig. E). In Fig. 3B, we plot the sequences obtained from this sampling onto two variables, their statistical energies *E*_GA_ and *E*_GB_, with the point corresponding to each sequence coloured by its committor *ϕ*_A_. This plot shows a clear separation of the sequences falling into GA and GB basins of attraction (according to committor value), with the variation of committor approximately correlated with the energy gap. However, while the gap is certainly a reasonable choice, in this case it is not optimal for separating the two folds as it is clearly not exactly orthogonal to the dividing surface [65] (Fig. 3B). An optimized version of the gap can be defined as 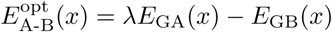 in which the optimal value of *λ* is chosen to maximize the correlation of the coordinate with the committor value (illustrated in S1 Text Fig. F). In Fig.3C and Fig. 3D we plot the free energy and position-dependent diffusion coefficients obtained from our MC simulations, for this coordinate. As a separate check of the quality of the optimized reaction coordinate, we compare the average value of the committor computed assuming 1D dynamics with the actual average determined over the sequences at each value of the coordinate. The similarity of the two curves, in Fig. 3A, demonstrates that 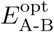(*x*) is indeed a good reaction coordinate for describing the dynamics [66] (in contrast, the agreement is not good using the unoptimized energy gap, as shown in S1 Text Fig. G).

### What is the Barrier to Fold Switching?

The barrier in the free energy on 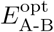 (*x*) is a measure of the difficulty of finding a path in sequence space between the two folds. For both move sets, there is a substantial barrier, ∼15*k*_B_*T* for all natural mutations and ∼30*k*_B_*T* when allowing only binary mutations (Fig. 3C). The higher barrier for binary mutations is anticipated due to the more restricted available sequences in that case. Although the dynamics we simulate is highly simplified as a model of protein evolution, the height of the free energy barrier, together with reasonable assumptions about the kinetic prefactor (based on replication error rates, population sizes and generation cycles), would suggest that this type of transition between folds is indeed a very rare event.

To further investigate the origin of the free energy barrier between the GA and GB basins in sequence space, we calculated the 2-dimensional free energy landscape projected onto *E*_GA_ and *E*_GB_ (Fig. 4C), based on umbrella sampling simulations in which all natural mutations were allowed. We see that the lowest free energy path from GA to GB does not follow a direct route, but rather an L-shaped path via a region where both *E*_GA_ and *E*_GB_ are large. In the context of our results on the correlation between protein stability and the statistical energies *E*_GA_ and *E*_GB_, the implication is that the most likely paths between folds go via unfolded, or unstable, states. We can obtain more insight into this by separating the free energy *F* (*E*_GA_, *E*_GB_) into its energetic *E*_comb_(*E*_GA_, *E*_GB_) and entropic *S*(*E*_GA_, *E*_GB_) = (*E*_comb_(*E*_GA_, *E*_GB_) −*F* (*E*_GA_, *E*_GB_))*/T* (Fig. 4B). Although the minimum energy path would clearly favour a direct transition from GA to GB, the very large contribution from sequence entropy favours a path through disordered states. In retrospect, this result seems obvious, given the vast size of unconstrained sequence space, relative to the size of the regions in which folds such as GA and GB are stable.

**Figure 4.**
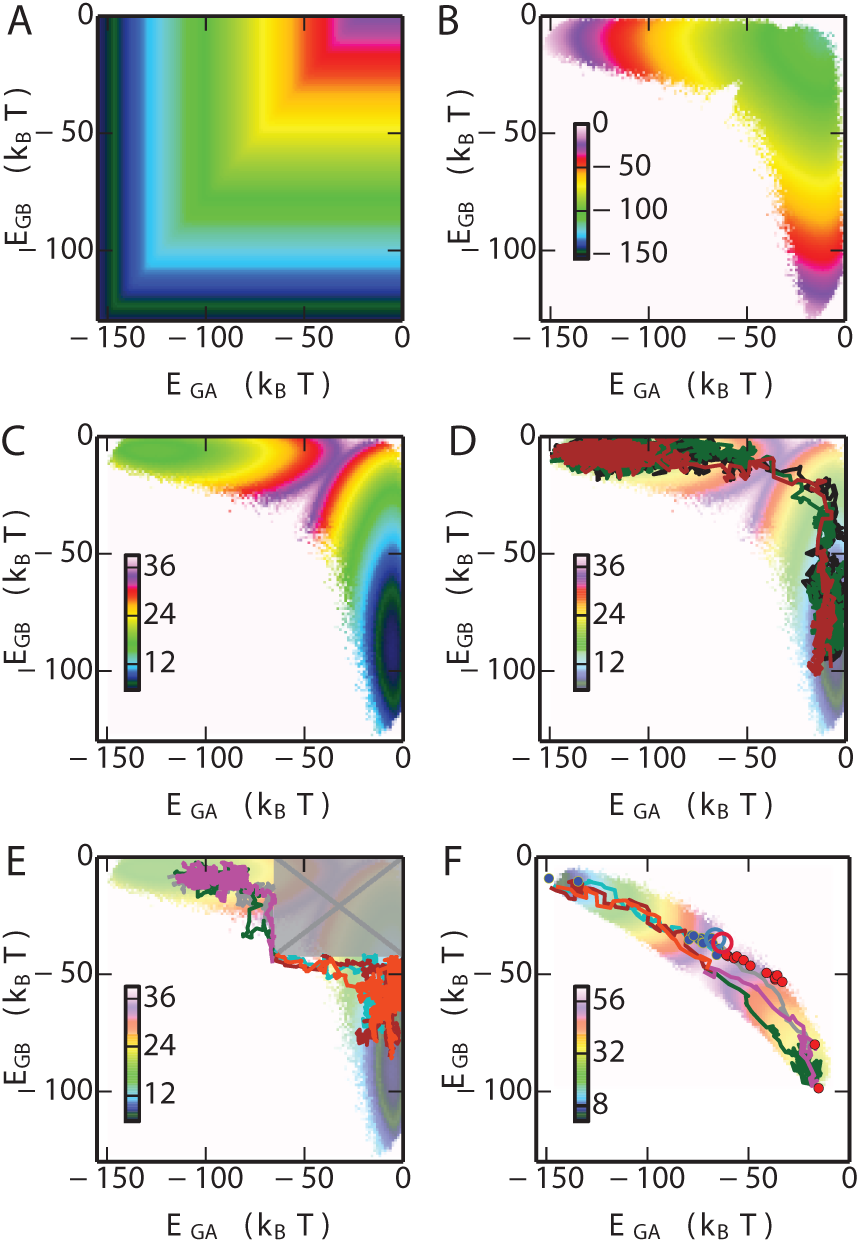
Fitness landscape. (A) Potential energy landscape of the combined model. (B) Contribution of entropy to free energy (times −1 for visualization). (C) 2D free energy landscape of the fold switch for natural mutation simulations. (D) Example of three transition paths from GA basin to the GB basin. Examples of transition paths (E) with stability constraints, and (F) using only “binary” mutations. The free energy surface in (F) is the one in which only binary mutations are allowed. All energies are in *k*_B_*T*.

### Transition Paths between Folds with and without Stability as an Evolutionary Pressure

In addition to calculating free energy surfaces from umbrella sampling, we have also determined directly examples of likely transition paths between the GA and GB folds. Since the free energy barrier between the two folds is very high (Fig. 3C), spontaneous transitions from one fold to another will rarely happen if using conventional sampling techniques. To obtain more statistics on the transitions, we used the transition path sampling technique (details in Methods), from which around 1000 transition paths on the fold bridge were obtained, a few of which are shown in Fig. 4D (with the remainder in S1 Text Fig. H). Consistent with the free energy surfaces, all paths go via sequences which have high values of *E*_GA_ and *E*_GB_, suggesting that in the absence of a constraint on protein stability, the most likely transitions from one fold to another involve sequences with lowered propensity for either fold. However, the average energies of the sequences in the transition region in Fig. 4D are still below zero, suggesting that some propensity for folding to GA and/or GB is retained even if the stability is low. We note that both experiments on GA/GB intermediates (see Fig. 2) [22, 56, 57], and simulations of simplified models [28, 34], have also suggested that loss of stability is invariably obtained as one approaches the bridge between folds.

Because in the cell unfolded chains would ordinarily be rapidly degraded, and because many proteins must be folded in order to function, the above scenario of fold conversion might be considered unrealistic. To avoid sampling sequences which are predicted to be unstable, we have also run transition-path sampling simulations in which the values of *E*_GA_ and *E*_GB_ are constrained to be below the boundaries separating stable and unstable sequences, −64.6 and −41.7 *k*_B_*T* for GA and GB respectively (Fig. 2C,D). The results of these runs, illustrated in Fig. 4E, show that there are still many possible paths allowed even with this restriction, consistent with the experimental finding of multiple stable bridge sequences. Interestingly, when only binary mutations are allowed (Fig. 4F), both the free energy surface and example transition paths suggest that the stability requirement is generally satisfied without having to be separately imposed. This follows from the much smaller sequence entropy contribution in this case; however, this restriction on sequence space also corresponds to a strong bias toward the target sequence. We note that the synthetic sequences on the fold bridge [22, 56, 57] (GA fold: blue, GB fold: red dot), also designed within the binary mutation space, fall within the bundle of transition paths sampled in this way (Fig. 4F). In addition Elber and co-workers have computationally designed, using the binary sequence space, a pair of sequences S1 and S2 which differ at one residue but are predicted to adopt the GB and GA folds respectively [67]. According to our model, S1 and S2, are shown as red and blue hollow circles in Fig. 4F, are very close to the fold interface. The *ϕ*_*A*_ of S2 and S1 are ∼1.0 and ∼0.87, respectively, suggesting that S2 has higher propensity to fold into GA topology than S1, consistent with the earlier prediction [67].

### Fold Bridge Sequences Are Likely to Be Intrinsically Disordered

What are the physical properties of the switch sequences (those with a committor *ϕ*_*A*_≃ 0.5) obtained from our simulations? A simple classification into sequences which favour globular structures and those which are more likely to be intrinsically disordered can be made on the basis of the mean net charge, *q*, and mean hydrophobicity, *h*. We have mapped the switch sequences obtained from our model using natural mutations onto these coordinates: Fig. 5A and B show, respectively, the results without and with a restraint on native state stability. On these plots, Uversky has determined that the line *q* = 2.785*h* − 1.151 [68] approximately separates IDP and globular sequences: by this criterion, 58% of the switch sequences without a restraint on native state stability fall into the IDP region, compared with only 26% when stability constraints are imposed. For reference, we have also calculated the *q* and *h* of experimentally well-characterized sequences from the IDP database DisProt [69] (Fig. 5C), with minimum disordered length > 4.) and the globular protein database Top8000 [70] (excluding those where regions of the sequence were not resolved in the structure). We find that 73% of the IDPs from DisProt and 8% of the globular proteins from the Top8000 are on the side of IDP as shown in Fig. 5C and D respectively.

**Figure 5.**
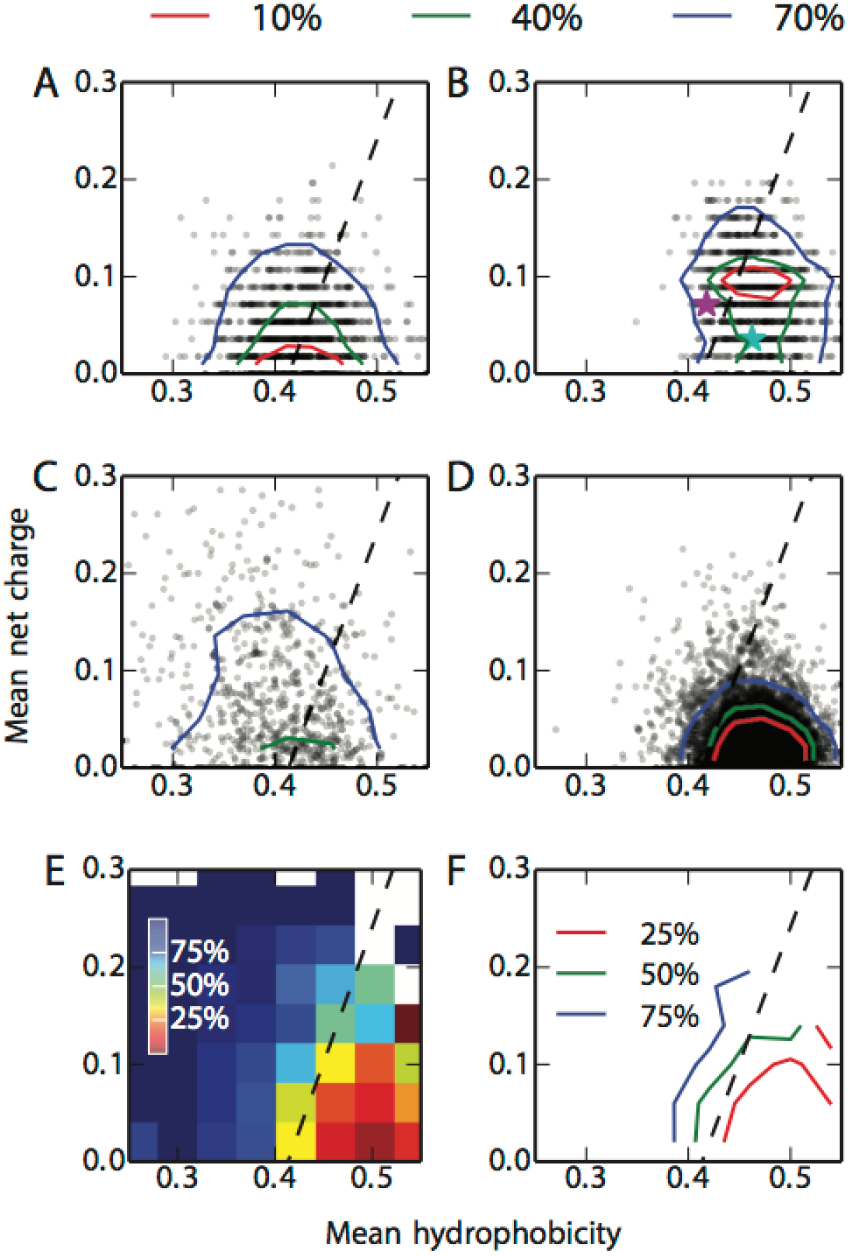
The Uversky plot divides proteins into folded globular and intrinsically disordered proteins based on their mean net change (*q*) and the mean hydrophobicity (*h*) [68]. In each plot, the dashed line represents the boundary between the two subsets described by Uversky [68]. We calculated the *q, h* of 10000 randomly selected transition sequences, defined as having *ϕ*_*A*_ within [0.49,0.51], from the simulations (A) without and (B) with stability constraints (one symbol for each sequence; probability density contours containing 10, 40 and 70 % of the data are also shown). The *q, h* of 694 known IDPs from the DisProt database [69] and 7957 globular proteins from the Top8000 database [70] are shown in (C) and (D) respectively. Sequences of GA and GB wild-type are shown with cyan and purple stars, respectively, in (B). (E) and (F) are respectively heat map and contour map representations of the IDP propensity *P* (IDP | *q, h*). The legends (%) represent the probability of being an IDP *P* (IDP | *q, h*) for each (*q, h*) combination.

It is clear from the reference data in Fig. 5C,D that the dashed line does not strictly separate IDPs and folded proteins. We have also employed a continuous descriptor, namely the conditional probability of being an IDP sequence for given values of *q* and *h, P* (IDP |*q, h*) (computed as described in Methods): this shows that indeed *P* (IDP |*q, h*) ≃50% near the previously determined dashed line (Fig. 5E,F). If we use a more conservative IDP descriptor, namely *P* (IDP |*q, h*) > 80%, we find 31 % and 6% of the switch sequences within this region without and with stability constraints, respectively. For comparison, of the simulated sequences from the two free energy basins of GA and GB, 1 % and 7 %, respectively, were in the IDP region.

Thus, by all measures considered, the switch sequences identified from our model without requiring the protein to be stable are enriched in sequences with a high propensity for disorder. This presents an alternative possibility to the scenario in which the folded state is constrained to be stable for all of the sequences bridging the two folds: the concern regarding possible aggregation or misfolding could be relieved by instead populating sequences with intrinsically disordered properties, namely low hydrophobicity and high net charge. Although unstable, these sequences would still have some propensity to fold to either GA or GB, as evidenced from their energies *E*_GA_ and *E*_GB_ being much below those for random sequences. The fact that it is much easier to find “bridge sequences” which are disordered than those that are folded may help to explain a growing catalog of IDPs which are able to fold into different structures upon binding with different ligands or other proteins [71], while such a property is very rarely observed for proteins which are independently stable. Whether evolution might take a similar route between folds is a matter of speculation, but an intriguing possibility nonetheless, considering the much greater probability of finding a path in this way.

### Key Residues Controlling Fold Switching

An obvious question concerning the switch sequences is whether there are any key regions of the sequence which are more important in determining the switch from the GA to GB folds…. Are there any common properties for the switch sequences? To identify the residues which play important roles for the fold switching, we analyzed the single site amino acid propensity during the fold switching when the stability constraints are imposed. The change of amino acid propensity from GA to GB sequence space at a given residue position can be indicated by 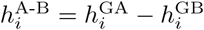, where *i* is the residue index, 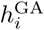 and 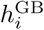 represent single-site propensities of GA and GB sequences respectively (Equation 1). The 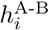 along the coordinate varies at different residues as shown in the Fig. 6A. At each residue, the overall changes of 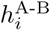 (indicated by *d*) and the rate of change in the transition region (indicated by *K*) where 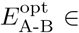 [−35.0, −33.0] (correspond to *ϕ*_*A*_ ∈ [0.1, 0.9]), are shown in the Fig. 6B and C respectively. The similarity between two probability distributions. At each residue position, to evaluate the similarity the probability distribution of the amino acid between the MSA of GA and GB, the Hellinger distance [72] (indicated by *δ*) is calculated as shown in Fig. 6D. Interestingly, we found that there is strong correlation between *δ* to either *d* or *K*. It suggests that the residues which play important roles in the fold switching are the ones that have the most distinct amino acid compositions in the MSAs of GA and GB. We also analyzed 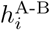 when no stability constraints are imposed, leading to a similar conclusion (S1 Text Fig. I).

**Figure 6.**
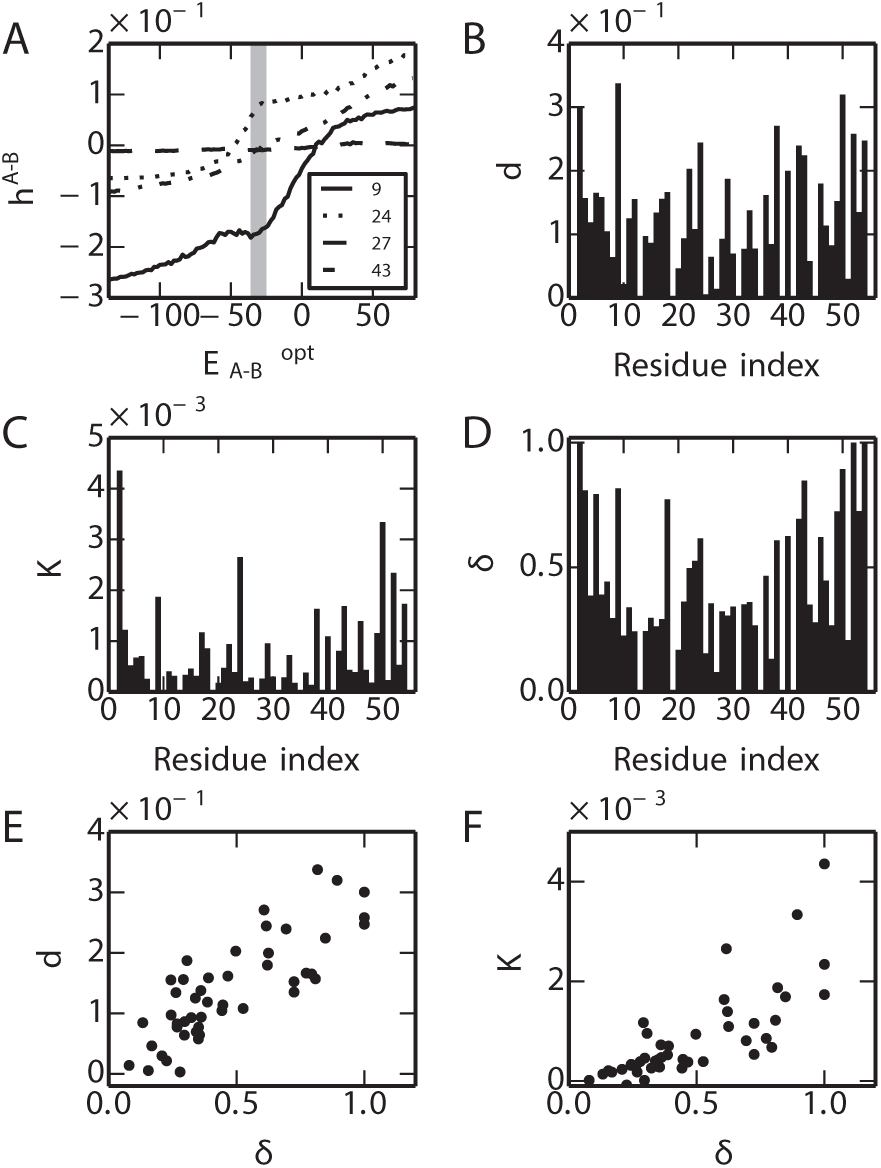
Single-site amino acid propensity changes in fold switching. (A) Examples of *h*^A-B^ for residues 9, 24, 27, 43. (B) Total change (*d*) of *h*^A-B^ from GA to GB. (C) Slope (K) at the transition region where it corresponds to *ϕ*_*A*_∈ [0.2,0.8]. The *δ* (D) and its correlation with *d* and *K* are show in (E) and (F).

## Discussion

We have generated a simple sequence-based model which successfully captures the propensity of all experimentally characterized sequences to fold into either the GA or GB structures, as well as separating the stable from unstable sequences. We have previously validated sequences designed using such models experimentally [39]. By using an ansatz inspired from energy landscapes in configuration space, we have combined the sequence-based fitness landscapes of the two folds to create a joint fitness function that can describe the propensity for both folds. We have used Monte Carlo dynamics to sample this joint fitness landscape in order to identify sequences with similar propensity for both folds. Such sequences could be considered as transition states on evolutionary paths between the two folds. More concretely, such sequences should be those most likely to switch folds apon single point mutation or binding to a cognate ligand.

Our results suggest that the number of possible bridge sequences at the interface of two folds is potentially very large (Fig. 4), even if the switch sequence is constrained to be stable (using the evolutionary Hamiltonian as a proxy for stability). Many of the bridge sequences generated from the simulation are predicted by the model to be of comparable or greater stability than the bridge sequences sampled in experiment [22, 56, 57]. The finding of multiple bridge sequences between folds may also be consistent with a recent analysis of the PDB suggesting that fold switching may be more common than previously thought [24].

Perhaps the most important conclusion from our study is that there are many more ways to find such fold-switch sequences which are unstable or have reduced stability. This is in qualitative accord with existing experimental and simulation studies [22, 28]. The reduction in stability may be expected to some extent based on the frustration between the sequence requirements of the two folds. Our study shows, however, that a second reason is the contribution from sequence entropy, which strongly favours a pathway via the more abundant low stability sequences. These low stability sequences tend to have properties usually associated with intrinsically disordered proteins (low hydrophobicity, higher charge content), raising the possibility that intrinsically disordered proteins may be able to function as bridges between protein folds in evolution [11, 73]. For example, there are several examples of IDPs that are known to fold to alternate structures when associating with different binding partners [71].

In future it will be interesting to apply this approach to design potential fold-switch sequences for this and other protein pairs which can be tested by experiment.

## Methods

### Multiple sequence alignments

The MSAs were generated with query sequences of GA (pdb code: 2FS1) [74] and GB (pdb code: 1PGA) [75] respectively, using the Jackhmmer method [76] (E-value cutoff: 10^−4^)) and the uniref90 database [77] (January, 2015). The MSAs contain 940 and 971 homologue sequences of GA and GB family respectively. Either *E*_GA_ or *E*_GB_ can successfully distinguish the sequences from different families (S1 Text Fig. A). For instance, the sequences of GA family have much lower energy than sequences of GB family under function *E*_GA_, and vice versa.

### Monte Carlo Sampling in the Space of Protein Sequence

The Metropolis-Hastings Monte Carlo method is employed here for the sampling guided by the combined energy potential *E*_comb_. In each Monte Carlo iteration, the amino acid of one random residue is perturbed by a flip, from one type of amino acid to another. All allowed types of amino acid at that position are attempted with equal probability. This takes the system from one sequence *x*′, with energy *E*_comb_(*x*), to a new sequence *x*′, with energy *E*_comb_′(*x*′). The move is accepted/rejected with acceptance probability

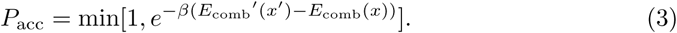

### First Passage Simulation and Transition Path Sampling in Sequence Space

Transition states are critical to understand the transitionary bridge connecting the GA and GB families. In the first passage simulation, the MC simulations start from random sequences and stop when it reach the boundary of the reaction coordinate which correspondeds to either free energy minimum of the fold. However, due to the high free energy barrier, full transitions from one fold to another happen very rarely by conventional sampling within reasonable timescale. Therefore, statistics around the transition region is very hard to obtain. We use transition path sampling [78] to overcome this bottleneck by starting simulations from amino acid sequences on the top of the free energy barrier. Simulations are running until it hit the boundary of either free energy basin.

### IDP Propensity Prediction

Given the mean net charge, *q*, and mean hydrophobicity, *h*, the probability of a sequence of being an IDP can be estimated from

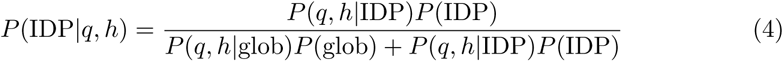

where *P* (IDP) and *P* (glob) are the estimated probabilities of IDP and globular proteins in nature, which are set to 30% and 70% respectively [3]. *P* (*q, h*| glob) represents the joint distribution of *q* and *h* in globular proteins and *P* (*q, h*| IDP) the distribution for IDPs. Here, *P* (*q, h*| IDP) is obtained from 694 IDP sequences from the DisProt database [69] (Fig. 5C) and the *P* (*q, h* | glob) is obtained from the 7957 Top8000 database of globular proteins [70] (Fig. 5D).

## Supporting Information

**S1 Text. Supporting text, tables and figures**. Procedure for verifying likelihood model via MC simulations (Text A). Wild type and designed amino acid sequences (Table A). Summary of stability and melting temperature of wild-type and designed sequences from previous experiments (Table B). Distribution of energies from Monte Carlo simulations with evolutionary Hamiltonian (Fig. A). Frequencies of amino acids at each position from Monte Carlo simulations with evolutionary Hamiltonian (Fig. B Correlation between evolutionary Hamiltonian and thermodynamic stability (Fig. C). Dependence of committor *ϕ*_*A*_ on *E*_comb_ (Fig. D). *E*_A-B_ of the designed sequences on the GA/GB fold interface (Fig. E). Determining optimal reaction coordinate from sequences with *ϕ*_*A*_ ≈ 1*/*2 (Fig. F). Quality of reaction coordinate assessed by comparison of true and calculated *ϕ*_*A*_ (Fig. G). Transition paths plotted on 2D fitness landscapes (Fig. H). Residues controlling fold switching (Fig. I).

## Acknowledgments

This work was supported by the Intramural Research Program of the National Institute of Diabetes and Digestive and Kidney Diseases of the National Institutes of Health. This study utilized the high-performance computational capabilities of the Biowulf Linux cluster at the National Institutes of Health, Bethesda, MD. (http://biowulf.nih.gov).

