## Supplemental text, tables and figures for "Exploring the Sequence Fitness Landscape of a Bridge Between Protein Folds"

### S1 Text

#### Text A

Before proceeding further, we tested whether sequences generated from the likelihood functions recapitulate the properties of the original MSA for each protein. To do this, we ran Metropolis Monte Carlo [1] simulations in sequence space, on either the $E_{\mathrm{GA}}$ or $E_{\mathrm{GB}}$ energy surface, as detailed in methods. As shown in SI Fig. S1, the energy distribution of the sequences from the MSA of GA or GB is consistent with the sequences generated by the Monte Carlo simulation using energy functions $E_{\mathrm{GA}}$ or $E_{\mathrm{GB}}$ scaled by a factor of 1.17 or 1.04 respectively. The rescaling of the energy allows us to recover the correct energy distributions by running simulations with each energy function at the same reduced temperature of 1.0. As a more detailed test, we compare the amino acid composition of the sequences from the MSA with those generated by the simulations (SI Fig.S4), finding good agreement

### Supporting Tables

Table A: Wild type and designed amino acid sequences.

| | |
|---|---|
| GA wild-type | MEAVDANSLAQAKEAAIKELKQYGIGDYYIKLINNAKTVEGVESLKNEILKALPTE |
| $^{63}GA_{MBP}$ | NGDKDANSLAEAKEKAIKELKIYGIGEHYIKLIENAKQVEAVESLKDEILKALPRF |
| $^{64}GA_{MBP}$ | NGDKDANSLAEAKEKAIKELKIYGIGEHYIKLIENAKQVAAVESLKDEILKALPRF |
| $^{68}GA_{MBP}$ | NGDKDANSLAEAKEKAIKDLKIYGIGEHYIKLIENAKQVAAVEDLKDEILKALPRF |
| $^{70}GA_{MBP}$ | NGDKDANSLAEAKEKAIKDLKIYGIGEHYIKLIEKAKQVAAVEDLKDEILKALPRF |
| $^{79}GA_{MBP}$ | NGDKGYNGLAEAKEKAIKDLKIYGIGEHYIKLIEKAKQVAAVEDLKDEILKAHDRF |
| $^{80}GA_{MBP}$ | NGDKGYNGLAEAKEKAIKDLKIYGIGEHYIKLIEKAKQVAAVEDLKDIILKAHDRF |
| GA30 | MEAVDANSLAQAKEAAIKELKQYGIGEKYIKLINNAKTVEGVWSLKNEILKALPTE |
| GA77 | TTYKLILNLKQAKEEAIKELVDAGIAEKYIKLIANAKTVEGVWTLKDEILKATVTE |
| GA77a | TTYKLILNLKQAKEEAIKELVDAAIAEKYIKLIANAKTVEGVWTLKDEILKATVTE |
| GA77b | TTYKLILNLKQAKEEAIKELVDAGTAEKYIKLIANAKTVEGVWTYKDEILKATVTE |
| GA77c | TTYKLILNLKQAKEEAIKELVDAGIAEKYFKLIANAKTVEGVWTLKDEILKATVTE |
| GA77d | TTYKLILNLKQAKEEAIKELVDAGIAEKYIKLIANAKTVEGVWTYKDEILKATVTE |
| GA77e | TTYKLILNLKQAKEEAIKELVDAGIAEKYIKLIANAKTVEGVWTLKDEIKKATVTE |
| GA77f | TTYKLILNLKQAKEEAIKELVDAGIAEKYIKLIANAKTVEGVWTLKDEILTATVTE |
| GA77g | TTYKLILNLKQAKEEAIKELVDAGIAEKYIKLIANAKTVEGVWTLKDEILKFTVTE |
| GA88 | TTYKLILNLKQAKEEAIKELVDAGIAEKYIKLIANAKTVEGVWTLKDEILTFTVTE |
| GA91 | TTYKLILNLKQAKEEAIKELVDAGTAEKYIKLIANAKTVEGVWTLKDEILTFTVTE |
| GA95 | TTYKLILNLKQAKEEAIKELVDAGTAEKYIKLIANAKTVEGVWTLKDEIKTFTVTE |
| GA98 | TTYKLILNLKQAKEEAIKELVDAGTAEKYFKLIANAKTVEGVWTLKDEIKTFTVTE |
| GB98-T25I | TTYKLILNLKQAKEEAIKELVDAGIAEKYFKLIANAKTVEGVWTYKDEIKTFTVTE |
| GB98-T25I-L20A | TTYKLILNLKQAKEEAIKEAVDAGIAEKYFKLIANAKTVEGVWTYKDEIKTFTVTE |
| GB98 | TTYKLILNLKQAKEEAIKELVDAGTAEKYFKLIANAKTVEGVWTYKDEIKTFTVTE |
| GB98a | TTYKLILNLKQAKEEAIKELVDAGTAEKYFKLYANAKTVEGVWTYKDEIKTFTVTE |
| GB95 | TTYKLILNLKQAKEEAIKEAVDAGTAEKYFKLIANAKTVEGVWTYKDEIKTFTVTE |
| GB91 | TTYKLILNLKQAKEEAIKEAVDAGTAEKYFKLYANAKTVEGVWTYKDEIKTFTVTE |
| GB88 | TTYKLILNLKQAKEEAITEAVDAGTAEKYFKLYANAKTVEGVWTYKDEIKTFTVTE |
| GB77g | TTYKLILNGKQLKEEAITEAVDAATAEKYFKLYANAKTVEGVWTYKDEIKTFTVTE |
| GB77f | TTYKLILNGKQLKEEAITEAVDAATAEKYFKLIANAKTVEGVWTYKDETKTFTVTE |
| GB77e | TTYKLILNGKQLKEEAITEAVDAAIAEKYFKLYANAKTVEGVWTYKDETKTFTVTE |
| GB77d | TTYKLILNGKQLKEEAITEAVDAGTAEKYFKLYANAKTVEGVWTYKDETKTFTVTE |
| GB77c | TTYKLILNGKQLKEEAITELVDAATAEKYFKLYANAKTVEGVWTYKDETKTFTVTE |
| GB77b | TTYKLILNGKQLKEEAIKEAVDAATAEKYFKLYANAKTVEGVWTYKDETKTFTVTE |
| GB77a | TTYKLILNLKQAKEEAITEAVDAATAEKYFKLYANAKTVEGVWTYKDETKTFTVTE |
| GB77 | TTYKLILNGKQLKEEAITEAVDAATAEKYFKLYANAKTVEGVWTYKDETKTFTVTE |
| GB30 | MTYKLILNGKTLKGETTTEAVDAATAEKYFKLYANDKTVEGEWTYDDATKTFTVTE |
| GB wild type | MTYKLILNGKTLKGETTTEAVDAATAEKVFKQYANDNGVDGEWTYDDATKTFTVTE |

Table B: Summary of stability and melting temperature of wild-type and designed sequences from previous experiments[2, 3, 4, 5].

| Protein | $\Delta G$ 25°C | $\Delta G$ 20°C | $T_m$/°C | Protein | $\Delta G$ 25°C | $\Delta G$ 20°C | $T_m$/°C |
|---|---|---|---|---|---|---|---|
| GAwt[2] | 6 | | 86.0 | GBwt[2] | 7 | | 87.5 |
| GA30[2] | 6 | | 86.0 | GB30[2] | 4.5 | | 65.0 |
| GA77[2] | 5 | 5 | 77.5 | GB77[2] | 4.0 | 5 | 62.4 |
| GA88[2] | 4 | | 69.4 | GB88[2] | 2 | | 57.5 |
| GA91[3] | | 4 | 61.5 | GB91[3] | | 3.1 | 49.3 |
| GA95[3] | | $\sim 3$ | 50.0 | GB95[3] | | $\sim 3$ | 48.7 |
| GA98[3] | | 1.5 | 37.0 | GB98[3] | | 2.0 | 35.0 |
| GA77d[3] | | 3.5 | 60.2 | GB98a[3] | | | 37.0 |
| GA77g[3] | | 4.7 | 75.3 | GB98-T25I[4] | | | 36.0 |
| GA77a[3] | | | 65.8 | GB77a[3] | | | 63.8 |
| GA77b[3] | | | 67.3 | GB77b[3] | | | 55.8 |
| GA77c[3] | | | 62.5 | GB77c[3] | | | 49.9 |
| GA77e[3] | | | 65.5 | GB77d[3] | | | 58.3 |
| GA77f[3] | | | 71.6 | GB77e[3] | | | 58.0 |
| $^{63}GA_{MBP}$ [5] | | | 80.0 | GB77f[3] | | | 61.9 |
| $^{64}GA_{MBP}$ [5] | | | 76.0 | GB98-T25I-L20A [4] | | | 46.0 |
| $^{68}GA_{MBP}$ [5] | | | 62.0 | | | | |
| $^{70}GA_{MBP}$ [5] | | | 63.0 | | | | |
| $^{79}GA_{MBP}$ [5] | | | 55.0 | | | | |
| $^{80}GA_{MBP}$ [5] | | | 53.0 | | | | |

### Supporting Figures

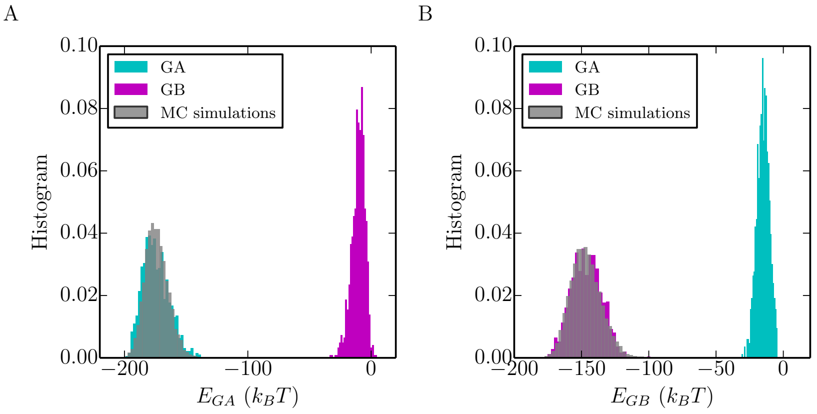


Fig. A: Distribution of $E_{\mathrm{GA}}$ (a) and $E_{\mathrm{GB}}$ (b) of homologous sequences from GA (cyan) and GB (purple) alignments. The distribution of $E_{\mathrm{GA}}$ and $E_{\mathrm{GB}}$ of the sequences sampled during Monte Carlo simulations are shown in grey. In the simulation, the energy functions of $E_{\mathrm{GA}}$ and $E_{\mathrm{GB}}$ have been scaled by a factor of 1.17 and 1.04 respectively, in order that the sampled distributions of energies match those from the original sequences.

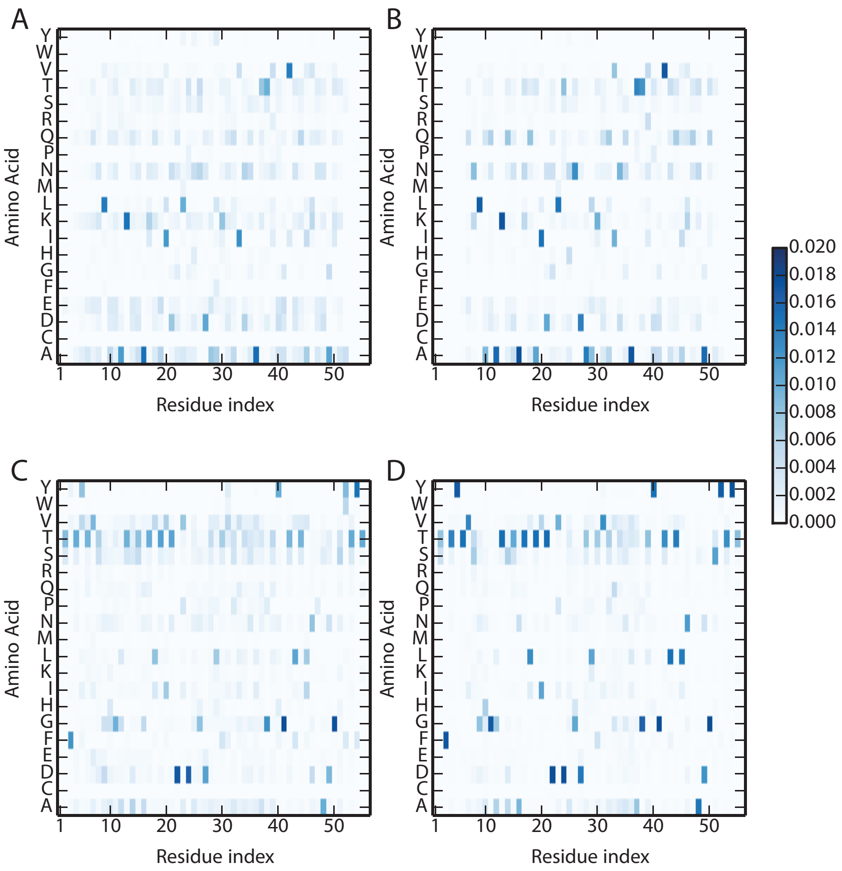


Fig. B: The single-site amino acid occupancies in the multiple sequence alignment of GA (A) are reproduced by the sequences generated by Monte Carlo simulations (B). Likewise, the single-site amino acid occupancies in the multiple sequence alignment of GB (C) are reproduced by sequences generated by MC simulations (D).

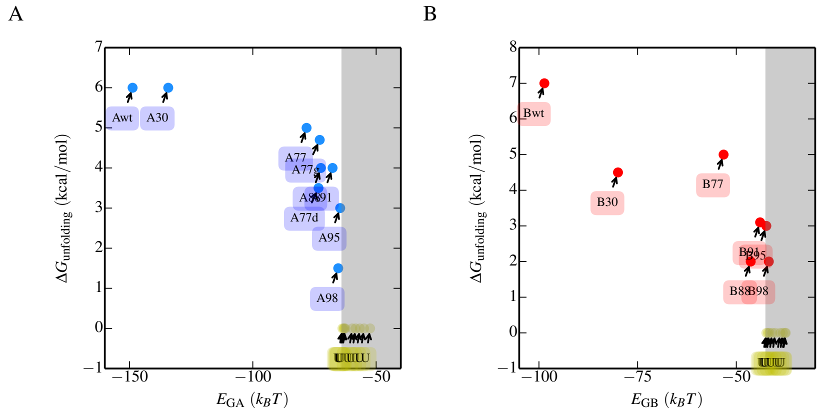


Fig. C: Evolutionary Hamiltonian and thermodynamic stability. The relation between stability $\Delta G_{\mathrm{unfolding}}$ and evolutionary Hamiltonian of GA and GB is shown in (A) and (B) respectively. $\Delta G_{\mathrm{unfolding}}$ are measured in the previous experiments [2, 3, 4].

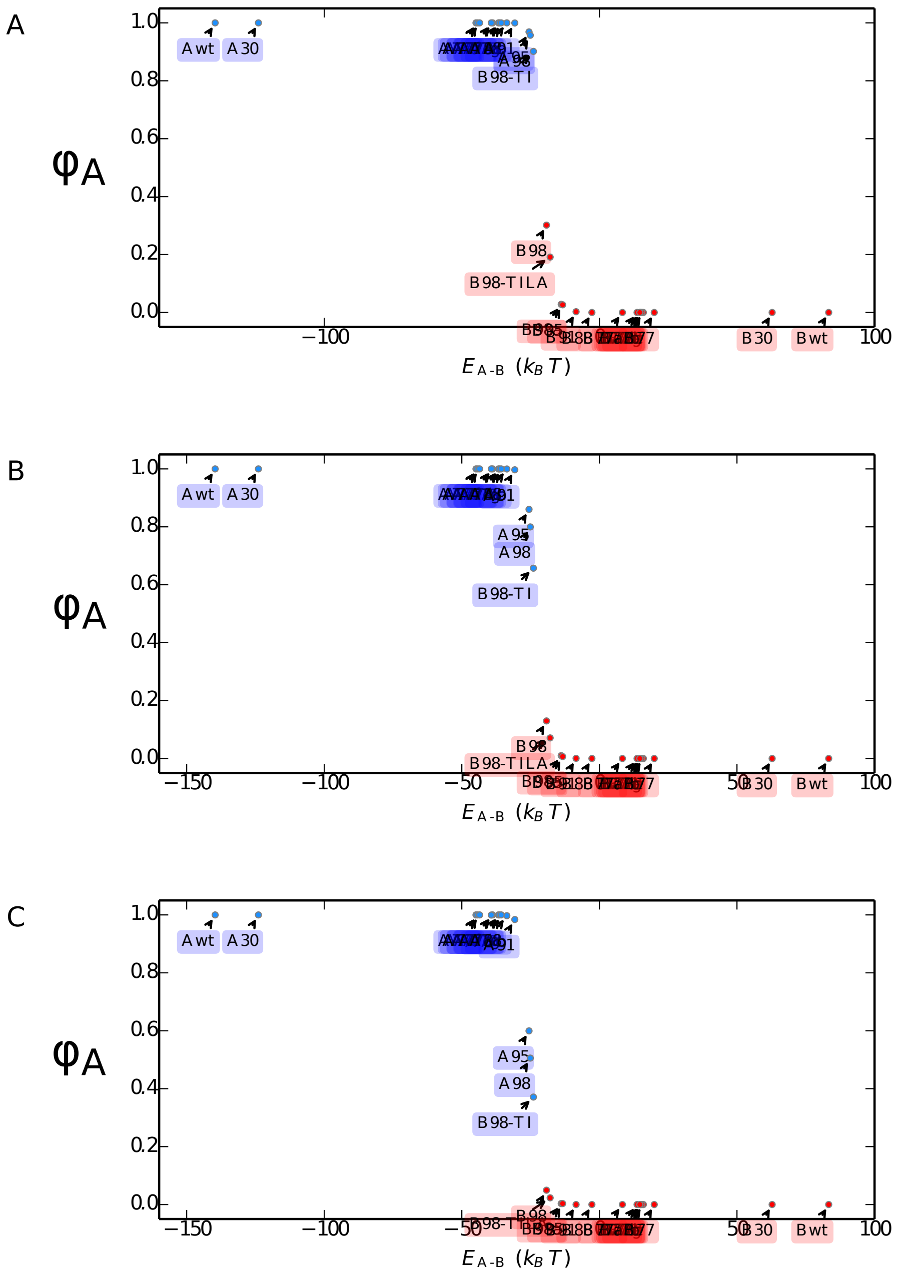


Fig. D: $\phi_A$ of mutations estimated by the first passage simulations. $\epsilon$ values of 21.0, 23.0, 25.0 are used in the figure A), B) and C) respectively.

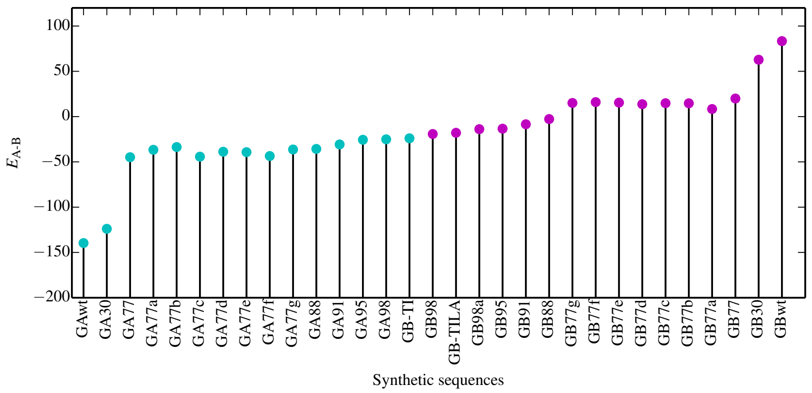


Fig. E: $E_{\text{A-B}}$ of the designed sequences on the GA/GB fold interface.

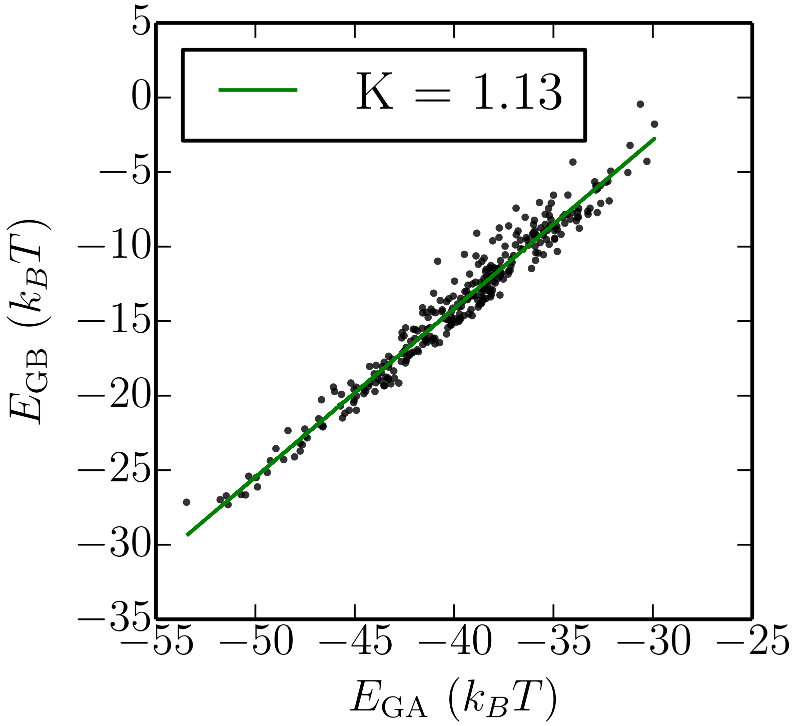


Fig. F: The designed sequences with $P_{\mathrm{fold}} \in [0.48, 0.52]$ are projected on the two coordinates $E_{\mathrm{GA}}$ and $E_{\mathrm{GB}}$. The least square fitting gives a straight line of slope K = 1.13.

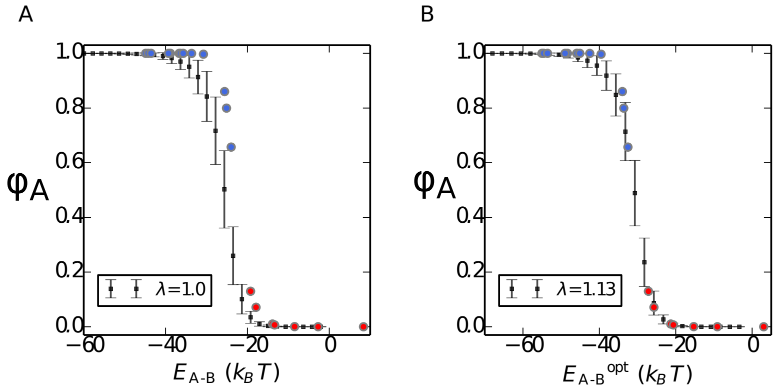


Fig. G: The mean and standard deviation of $\phi_A$ is calculated along the reaction acoordinate $E_{\text{A-B}}$ with $\lambda = 1.0$ and the optimized one $E^{\text{opt}}_{\text{A-B}}$ with $\lambda$ = 1.15.

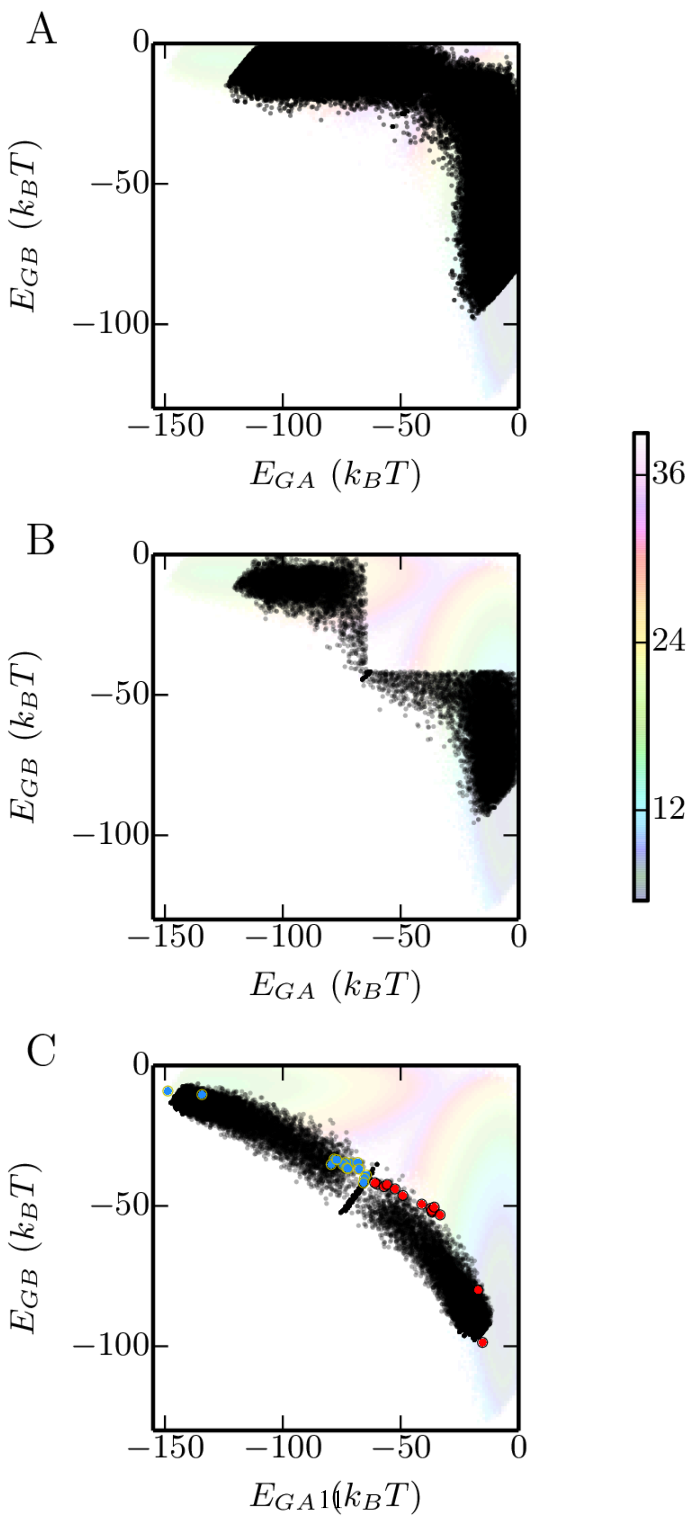


Fig. H: The transition path sequences are plotted in black dots for the case of natural mutation (A), natural mutations with stability constraints (B), and binary mutations (C). Red and blue dots are experimental mutations with GA and GB folds respectively.

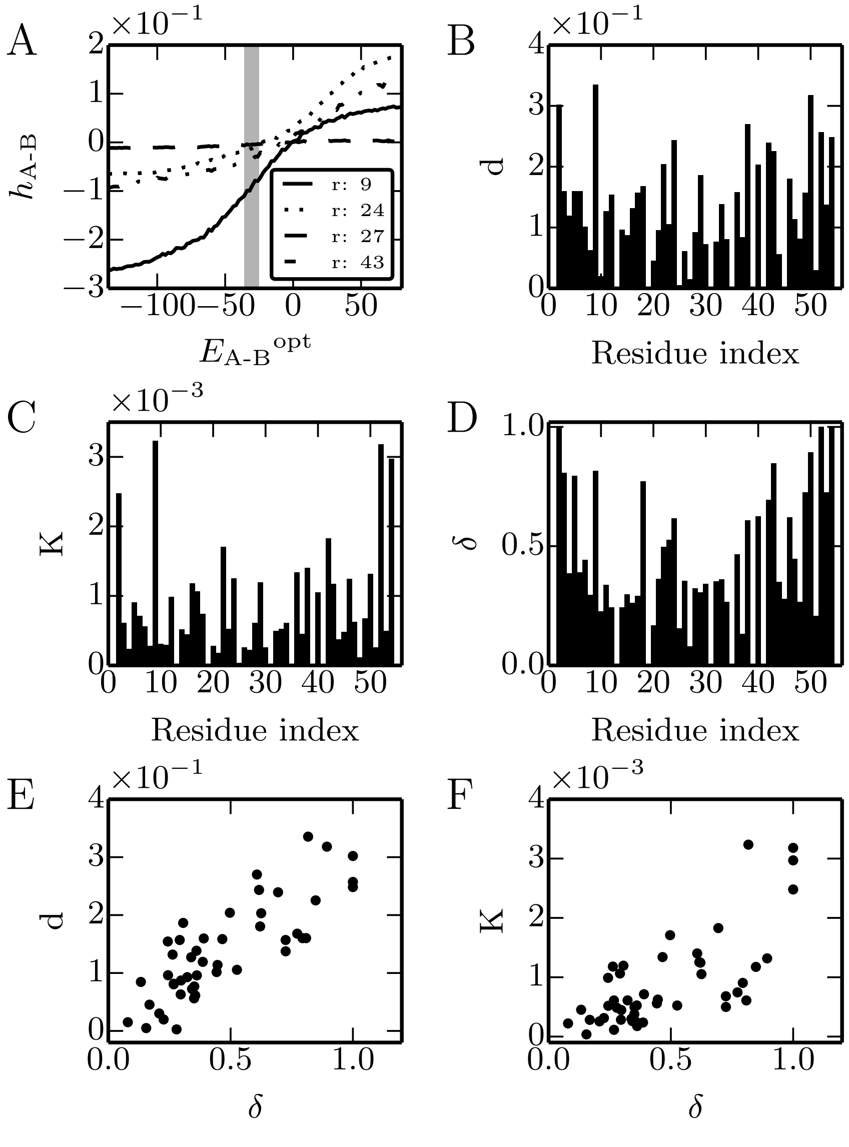


Fig. I: Natural mutations. (A) examples of Hb-Ha for residues 20, 25, 45. (B) Total change (d) of Hb-Ha from GA to GB. (C) Slope K at transition. (D) The difference ($\delta$) in residue propensity between GA and GB homologs. (E) Correlation of d with $\delta$. (F) Correlation of slope with propensity.
